# RecA is a reliable marker for bacterial taxonomy, even in the Candidate Phyla Radiation

**DOI:** 10.1101/2024.06.21.600076

**Authors:** Lodovico Sterzi, Simona Panelli, Clara Bonaiti, Stella Papaleo, Giorgia Bettoni, Enza D’Auria, Gianvincenzo Zuccotti, Francesco Comandatore

## Abstract

Culture-independent approaches are commonly used to characterise the taxonomic composition of bacterial communities. Among these approaches, the amplicon-based metagenomics relies on specific genetic markers, such as the 16S rRNA gene, while the shotgun metagenomics annotates the whole bacterial DNA. Despite the 16S being the gold standard marker, studies highlighted its inefficiency in characterising and quantifying divergent bacterial groups such as the Candidate Phyla Radiation. On the other hand, shotgun metagenomics is highly informative and accurate but it is more expensive and requires computational resources and time. In this study, we propose RecA as a pan-bacterial genetic marker, particularly suitable for the Candidate Phyla Radiation. Indeed, we found that applying a Random Forest machine learning model on RecA amino acid sequences provides an accurate and fast taxonomic annotation across the whole bacterial tree of life. Ultimately, we produced Forestax, a tool for the characterisation and quantification of bacterial communities in metagenomics data, on the basis of RecA sequences. The analyses showed that RecA-based metagenomics has a taxonomic accuracy comparable to other multi-gene approaches, reinforcing RecA as a powerful marker for taxonomic annotation in bacteria. In perspective, RecA could be considered as a broad-spectrum marker for amplicon-based studies to overcome the limits of 16S rRNA.

## Introduction

The accurate taxonomic identification of bacteria in a biological sample represents a long-standing, major challenge in microbiology. With the advent of high-throughput sequencing techniques, microbial communities have been commonly described via culture-independent procedures. These procedures involve either sequencing a specific marker sequence (amplicon metagenomics) or all genomic content (shotgun metagenomics) in a biological sample^1–3^.

In amplicon metagenomics, the bacterial taxonomic composition is determined by analysing a marker sequence, with the small subunit of the ribosomal RNA (16S rRNA) gene being the most frequently used marker^4–6^. First proposed by Carl Woese in 1977, the 16S rRNA gene (16S) presents evolutionary and genetic features which make it an ideal marker: it is present in all bacteria, it is seldom horizontally transferred and it is composed by a combination of conserved and variable regions^7^. In amplicon metagenomics, the 16S in a given DNA sample is amplified, sequenced and taxonomically annotated using bioinformatic pipelines. This technique is robust, standardised and cost-effective, but limits are present. It has a limited discriminatory power, in particular in less explored parts of the tree of life, such as the Candidate Phyla Radiation (CPR) monophylum^8,9^. Indeed, CPR bacteria exhibit notably divergent 16S sequences, rendering universal 16S rRNA primers less effective and remaining largely unseen by amplicon-based studies^10–12^. Another limit is that the 16S rRNA gene can be often found in multiple copies in several bacterial species, severely affecting the relative abundance estimates^13^. During the last ten years, metagenomics based on the sequencing of the whole bacterial DNA sample (i.e. shotgun metagenomics) has become increasingly popular to describe complex bacterial communities. Several methods have been developed to delineate the taxonomic composition within a microbial community starting from genomic data^14^. A method is to assemble raw reads (i.e. the metagenome) and taxonomically assign the obtained contigs against a reference sequence database. Since this approach is a computational burden, softwares which performs the taxonomic annotation directly from raw reads has been proposed^15–19^. The main advantage of shotgun metagenomics is that it is not affected by 16S-specific biases like the presence of multiple copies and the primer specificity. On the other hand, it is less accessible than amplicon metagenomics, requiring high bioinformatic skills, time and computational power.

At the state of the art, a huge amount of shotgun metagenomics datasets are available in public databases, but high-throughput analysis remains computationally challenging. Using a single marker gene for metagenomics analysis could reduce the required computational power but, unfortunately, the 16S rRNA gene is not efficiently assembled in shotgun metagenomics data.

In this study we analysed thousands of bacterial genomes to identify a novel marker gene, alternative to 16S, for bacterial taxonomy assignment, with a particular focus on Candidate Phyla Radiation. The results highlight that RecA is a very promising target. Thus, we developed Forestax, a scalable, user-friendly highly efficient machine learning-based tool for bacterial taxonomy assignment in very large datasets, on the basis of the RecA sequence. We can not exclude that the same approach/tool could be used in the future in RecA-based amplicon metagenomics.

## Methods

### Identification of a protein marker for the Candidate Phyla Radiation

#### Dataset reconstruction

The global Candidate Phyla Radiation (CPR) genomic dataset was composed by all “Patescibacteria’’ genome assemblies present in both Genome Taxonomy Database (GTDB)^20^ and Bacterial And Viral Bioinformatics Resource Center (BV-BRC) database^21^. Assembly statistics were obtained with assembly-stats (https://github.com/sanger-pathogens/assembly-stats) and full 16S sequences were extracted using Barrnap (https://github.com/tseemann/barrnap). Gene prediction was performed using Prodigal ^20,22^, adjusting the genetic code from standard code 11 to code 25 for Gracilibacteria and Absconditabacteria.

#### Identification of single-copy core protein clusters

To find a single-copy core genetic marker, the global CPR genomic dataset was refined: CPR genome assemblies without gaps, without unidentified bases and with a fully-assembled 16S sequence were retained in the refined CPR genomic dataset.

The orthology analysis was performed as follows. First, DIAMOND^23^ was used with a sensitive approach (--sensitive) for an all-against-all comparison of protein sequences in the refined CPR genomic dataset. Then, a graph of protein sequences was composed where two sequences were connected only if the comparison revealed a hit with coverage (length of the hit / length of the query sequence) between 0.8 and 1.1 and for sequence identity above 70. Then, R library *igraph* was used to decompose the graph in separate protein clusters. Protein clusters present in at least 90% (409/454) of genomes were identified.

Misassembled and/or partial protein sequences were removed by filtering for sequence length outliers. Then, retained sequences were annotated against the COG^24^ database to identify protein clusters which corresponded to single proteins. Protein clusters in which at least 98% of sequences were assigned the same COG number were retained and, if necessary, filtered from other low-frequency (<= 2%) COGs.

After the filtering steps, the presence of protein clusters in the strains was re-evaluated. Protein clusters which were present in at least 90% of genomes (409/454) and in multiple copies in less than 1% of genomes (5/454) were retained.

#### Evaluation of protein markers taxonomical discrimination

Sequences in the retained protein clusters were aligned using Muscle^25^ and subjected to Maximum Likelihood (ML) phylogenetic analysis using FastTree MP^25,26^, using the general time reversible (GTR) model. Phylogenetic trees were manually inspected and protein clusters which could discriminate the main CPR groups within the refined CPR genomic dataset were considered good candidates.

The proteins were searched by DIAMOND against the global CPR genomic dataset (see the “Dataset reconstruction” section) and the additional protein sequences were added to the clusters.

To include similar protein representatives from non-CPR bacteria, a dataset of non-CPR bacteria was downloaded from GTDB. Gene calling was performed by Prodigal and their protein content was screened for the presence of the candidate protein clusters using DIAMOND (with the same thresholds used above).

Finally, all sequences from the candidate protein clusters were realigned and re-subjected to the phylogenetic analysis performed above. The best candidate was chosen by manual inspection of the phylogenetic tree.

### Random forest model for taxonomic assignment

#### Pan-bacterial RecA database construction

All available bacterial genome assemblies were downloaded from GTDB on September 14, 2023 and gene prediction was performed using Prodigal, adjusting the genetic code from standard code 11 to code 25 for Gracilibacteria and Absconditabacteria. Then, protein sequences from each genome were compared via DIAMOND (--evalue 0.00001, --max-target-seqs 5) to RecA protein sequences from the COG database (COG0468). Sequences with mean query coverage and subject coverage between 0.5 and 1.5 were retained to create a pan-bacterial RecA protein dataset. Sequences with query coverage and subject coverage between 0.95 and 1.05 were marked as complete.

#### Random Forest Taxa Assignment Models

The RecA sequences were translated to the compressed Dayhoff(6) alphabet and kmers of length 7 were counted using the *kmer* R library.

Then, six Random Forest prediction models (one for each GTDB taxonomic level) were built for taxonomy assignment of the RecA sequences.

For each model, complete RecA sequences were randomly split in a training dataset and a testing dataset with a 7:3 ratio, stratifying on the taxonomic level. Taxa with less than 10 representatives were discarded. Probability forests were built on the kmers using the “ranger” R library. Each forest was composed of 1000 trees, with 280 independent variables (kmers) sampled in each tree node.

#### Random Forest Reliability Models

To account for the uncertainty of the taxonomic assignment, six additional Random Forest models (one for each GTDB taxonomic level) were built to give a reliability score to each taxonomic assignment.

To develop these models, the testing datasets of the taxonomy assignment models were sub-split in training and testing datasets (again with a 7:3 ratio, as above).

The probability forests were built on three independent variables: query coverage, subject coverage and probability of the taxonomic assignment. Each forest was composed of 1000 trees, with one independent variable sampled in each tree node.

#### Forestax overview and evaluation on mock community samples

Forestax was developed in the following way: the tool takes as inputs either paired-end raw reads, (meta)genomic assemblies or protein sequences. It extract putative RecA proteins and provide a taxonomic assignment using the Taxa Assignment Models and Reliability Models.

If the inputs are raw reads, the tool extracts reads belonging to the RecA gene using Bowtie against an index composed by the pan-bacterial RecA database (see “Pan-bacterial RecA database construction”) and assembles the reads using SPAdes^27^. The tool then proceeds to call the coding sequences using Prodigal and identifies putative RecA sequences by DIAMOND against a refined RecA COG database. RecA protein sequences are translated to the compressed Dayhoff(6) alphabet and length 7 k-mers are counted. The Taxa Assignment Models and Reliability Models take the matrix as input and predict the taxonomy for each RecA sequence. Assignments with a Reliability score < 50/100 are marked as unclassified.

The tool can also operate starting from assembled nucleotide sequences and protein sequences.

Relative abundances can be inferred from raw reads using the mean read coverage obtained by Bowtie2^28^ and Samtools^29^.

To evaluate the tool performance on mock bacterial communities, the raw reads of 22 mock community samples used in Valencia et al were downloaded from NCBI. The tool Forestax was used with the following parameters:

*-itype reads -o output_folder -r1 reads_fw.fastq.gz -r2 reads_rv.fastq.gz -cpu 50 –q -tax phylum,class,order,family,genus,species*.

The performance of the tool was evaluated at genus and species level using the same metrics as Valencia et al^30^: Sensitivity, False Positive Relative Abundance (FPRA) and Aitchison Distance.

- Sensitivity was calculated as: (number of correctly identified species / the number of expected species)*100.
- FPRA was calculated as: (abundance of false positive species / total abundance)*100.
- Aitchison Distance was computed in R with function aDist of library “robCompositions”. To deal with zeroes, the multiplicative replacement method with *delta* 0.001 was used.

As done in Valencia et al, for replicate communities, mean and standard deviation values were computed. The values obtained for Forestax were merged to the output table obtained by Valencia et al. for the other pipelines and plotted in R.

### Statistics and Reproducibility

This study was conducted on bacterial genome assemblies available on NCBI and included in the GTDB/BV-BRC databases.

Random Forest models were developed using the “ranger” R library and dividing the data in a training dataset and a testing dataset, stratifying on taxonomic groups.

Data analysis was performed using R. Plots, box plots, histograms and bar charts were produced using the *ggplot* and *ggpubr* R libraries.

The non-parametric Mann-Whitney U test was used to compare the distribution of continuous variables in independent samples.

## Results

### Protein marker selection

To identify a strong protein marker for the profiling of bacteria belonging to the Candidate Phyla Radiation (CPR), we created a global CPR dataset of 3971 genomes marked as “Patescibacteria” in the Genome Taxonomy Database (GTDB) (Supplementary Table S1).

We performed a custom orthology analysis on a refined CPR dataset of 454 high-quality genomes, combining a very strict sequence clustering analysis, COG annotation and phylogenetic analysis to extract single-copy core orthologous groups without close paralogs (see the Methods section). The protein clustering analysis identified nine suitable protein clusters and COG annotation revealed that these protein clusters corresponded to proteins involved in core cellular functions (Supplementary Table S2).

We excluded protein clusters which showed clear signals of paralogy and we performed a phylogenetic analysis to identify a target which could well discriminate the main lineages in the CPR. Two protein clusters were considered suitable (Cluster 2 and Cluster 19, Figure S1) and they were searched in the global CPR dataset. Furthermore, we composed an additional dataset of 1251 bacterial genomes belonging to 145 non-CPR orders and searched the protein clusters within these genomes, to test whether the potential protein markers could distinguish CPR bacteria from other groups. As shown in Figure 1, the phylogenetic tree of Cluster 2, annotated as RecA protein, was strongly coherent with the CPR evolutionary history and distinguished the main lineages within the group. Thus, the RecA protein was chosen as the best candidate.

**Figure 1.**
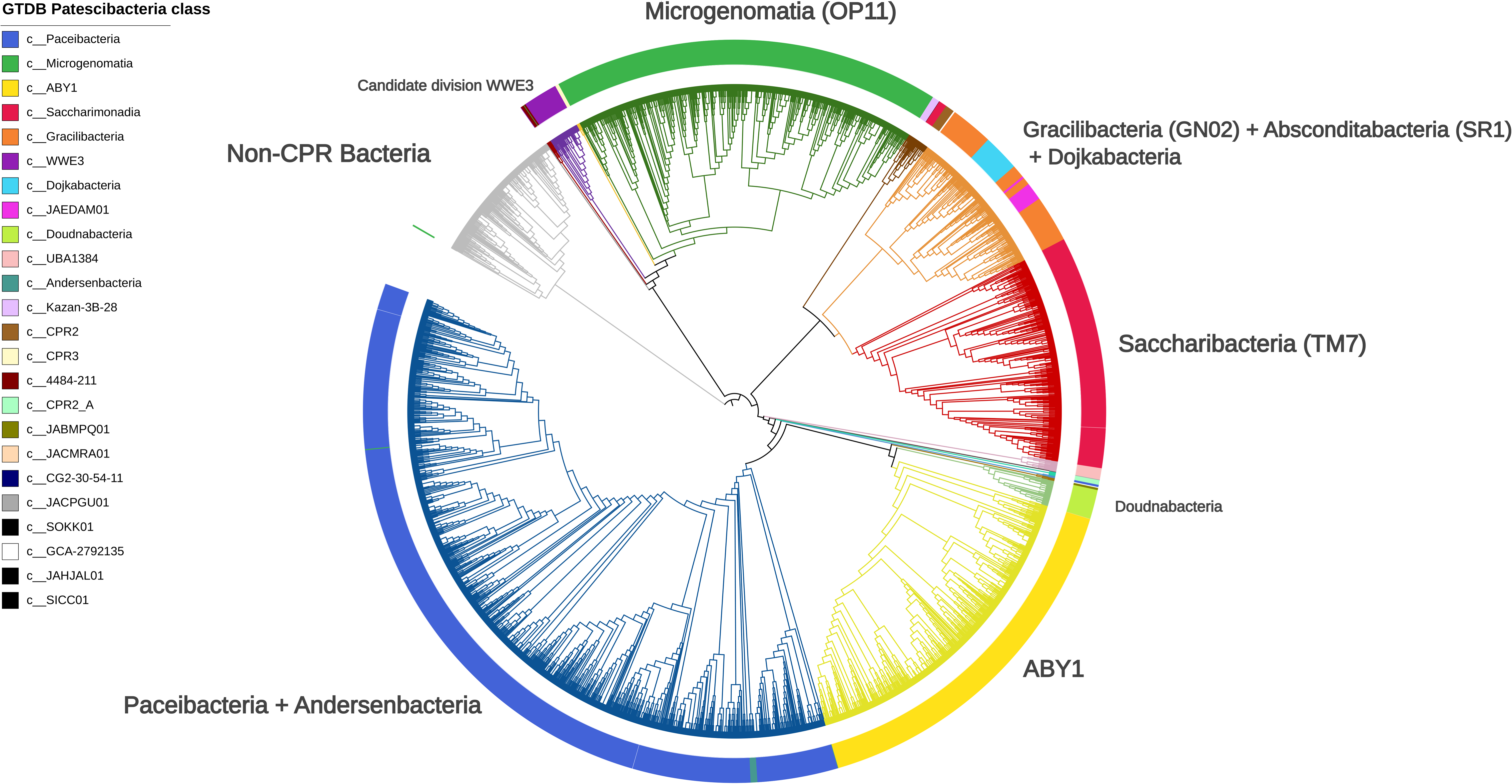
ML phylogenetic tree of protein Cluster 2, annotated as RecA protein. The coloured ring around the tree indicates the taxonomic class origin of the RecA sequence from the Candidate Phyla Radiation, according to the Genome Taxonomy Database. Branches are coloured according to the main RecA lineages highlighted by the tree topology.

To test if this marker could be used for a broader taxonomic profiling in the bacterial domain, we created a pan-bacterial RecA protein dataset.

First, we extracted the RecA protein from a dataset of 309,845 bacterial genome assemblies found in the Genome Taxonomy Database (GTDB). We found a total of 316,029 putative RecA proteins: only 8,386/309,845 genomes (2.7%) did not have a RecA sequence, whereas 287,835/309,845 genomes (92.9%) had one putative RecA sequence and 13,624/309,845 genomes (4.4%) had more than one sequence. On the basis of the hit coverage, we distinguished the protein sequences into 260,504 complete sequences and 55,525 partial/misassembled sequences (see Methods).

Across bacterial diversity, the RecA protein was found in 58,228/61,614 species (94.5%). This frequency increases to 11,681/11,740 species (99.5%) when excluding species represented by less than three genomes. Among the 59 species lacking RecA, we highlight the presence of obligate endosymbionts with reduced genomes such as *Buchnera aphidicola* and *Sulcia muelleri*.

All information on the RecA pan-bacterial dataset can be found in Supplementary Table S3.

### Models training and validation

To develop a taxonomy classification model from the pan-bacterial RecA protein dataset, we trained and tested six Random Forest models, one for each GTDB taxonomic level, on 260,504 complete RecA sequences. From now on, we will refer to these random forest models as the Taxa Assignment Models.

The Taxa Assignment Models analyse the kmer composition of the RecA protein sequence and return a probability score matrix of its bacterial taxon (see Methods). The taxon with the highest score is assigned.

First, we evaluated how well the models performed on ∼70,000 complete RecA sequences. For each taxonomic level, we annotate two metrics: the percentage of correctly assigned sequences and the percentage of well assigned taxa. We considered a taxon well assigned if at least 90% of sequences in the taxa were correctly assigned. As visualised in Figure 2a, the models were able to assign the correct taxon down to the species level for 51,199/56,895 sequences (90%), with 1,138/1,583 species (72%) well assigned evenly across 132 bacterial orders. As shown in Figure 2b and Table 1, the models assigned the correct genus for 70,556/71,283 sequences (99%), with 1,332/1,438 genera (93%) well assigned. At higher taxonomic levels, the models assigned the correct taxon to approximately 99.8% of the sequences, while the number of well assigned taxa was 632/673 (94%) at the family level, 326/367 at the order level (88.9%), 111/149 at the class level (74.5%) and 50/72 at the phylum level (69.5%). Within the Candidate Phyla Radiation, the assignment models could identify as Patescibacteria (the GTDB phylum corresponding to the CPR) 653/656 RecA sequences (99.5%). The main lineages in the CPR were distinguished correctly at the class for 648/655 sequences (98.9%), with 8/10 classes (80%) well predicted, and at the correct order for 518/526 sequences (98.5%), with 30/31 orders well assigned.

**Figure 2.**
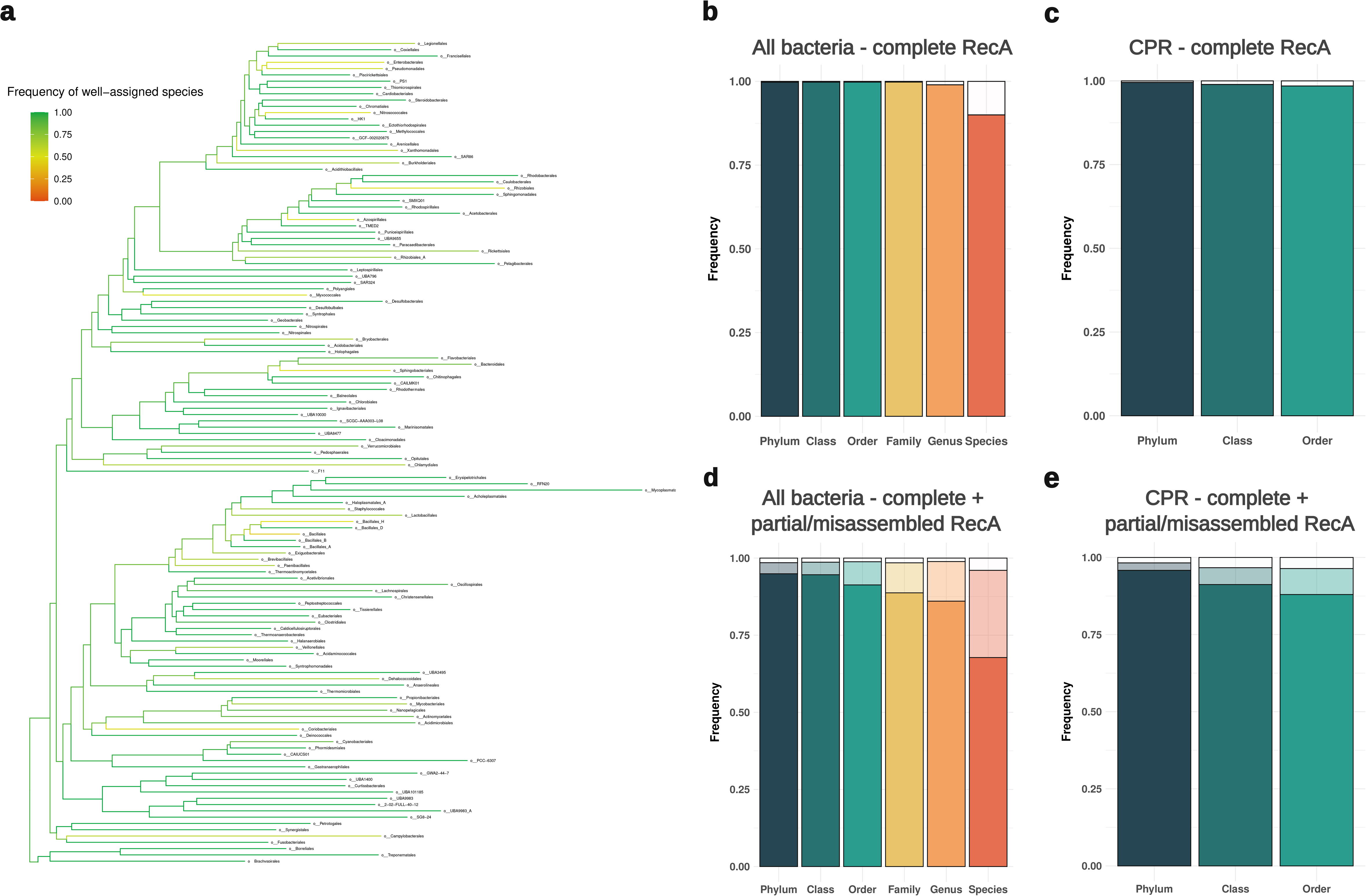
a) Phylogenetic tree (obtained from the Genome Taxonomy Database) with branches coloured on the basis of the frequency of well-assigned species (>90% of sequences assigned to the correct species) across 132 bacterial orders. Taxonomic assignments were performed on complete RecA sequences. Internal nodes are coloured by the mean value of the orders within the node. b) Bar plots showing frequencies of complete RecA sequences assigned to the correct taxon at each taxonomic level for all bacteria. c) Bar plots showing frequencies of complete RecA sequences assigned to the correct taxon at each taxonomic level for groups within the Candidate Phyla Radiation (CPR). d) Bar plots showing frequencies of complete and partial/misassembled RecA sequences assigned to the correct taxon at each taxonomic level for all bacteria. e) Bar plots showing frequencies of complete and partial/misassembled RecA sequences assigned to the correct taxon at each taxonomic level for groups within the Candidate Phyla Radiation (CPR). The coloured portion of the bar plot indicates correct taxonomic assignments, while the transparent portion of the bar plot indicates unclassified sequences (i.e. sequences which could not be confidently assigned to a taxon). Since lower taxonomic levels within the CPR are not well-defined, assignments of RecA sequences belonging to CPR bacteria were analysed to the order level.

**Table 1.** Table summarising the percentage of correct taxonomic annotations obtained by the machine learning models on RecA protein sequences.

The percentage of RecA sequences assigned to the correct taxon was related to the number of sequences available for that taxon, indicating a higher model/marker accuracy for more represented bacteria (Figure S2).

### Training and testing reliability models for partial/misassembled RecA sequences

As stated above, the inclusion in the dataset of partial/misassembled RecA sequences (i.e.sequences annotated as RecA against the COG database with unusually high or low coverage) significantly increases the number of single-copy RecA sequences found in the pan-bacterial genome dataset (Figure S3a). Thus, the models were also tested on a dataset of misassembled/partial RecA sequences. As shown in Figure S3b, the inclusion of these sequences in the analysis significantly reduced the assignment accuracy at all taxonomic levels. At the genus level, only 25,280/39,512 (64%) of partial sequences were assigned to the correct genus and only 2,940/21,682 (13.56%) of the partial sequences were assigned correctly at the species level. Overall, model accuracy was below 90% at all taxonomic levels: all the results for complete and partial sequences are summarised in Table 1 and Supplementary Table S4.

We analysed the model results and determined that wrong assignments could be distinguished from correct assignments on the basis of the protein coverage values and on the basis of the probability given to the assignment by the Taxa Assignment Model (Mann-Whitney U Test, p < 0.01, Figure S4). Thus, we trained six additional random forest models (one for each taxonomic level) to assess the reliability of the Taxa Assignment Models on the basis of these parameters. From now on, we will refer to these models as the Reliability Models. The Reliability Models were trained and tested on datasets including complete sequences and partial/misassembled sequences, providing to each assignment a score between 0 and 100. Taxonomic assignments with a low probability of being correct (i.e. score < 50/100) were labelled as unclassified. As shown in Figure 2c, at the species level the Reliability model was imprecise (i.e. a wrong assignment considered likely/a right assignment considered unlikely) for 939/23,574 sequences (4%) and precise for 22,635/23,574 sequences (96%): 6,662/23,574 sequences (28.3%) are recognised as unclassified and 15,973/23,574 sequences (67%) are correctly assigned. At the genus level, the model is imprecise for 372/32,867 sequences (1.1%) and it is precise for 32,495/32,867 sequences (98.9%): 4,270/32,867 sequences (12.9%) are recognised as unclassified and 28,597/32,867 sequences (86%) are correctly assigned.

Within the Candidate Phyla Radiation, at the class level the model is imprecise for 28/843 sequences (3,3%) and at is precise for 815/843 sequences (96.7%): 46/843 sequences (5.5%) are unclassified and 769/843 are correctly assigned (91.2%). At the order level, the model is imprecise for 26/724 sequences (3.6%) and it is precise for 698/724 sequences (96.4%): 61/724 sequences (8.4%) are unclassified and 637/724 (88.0%) are correctly assigned.

Overall, the inclusion of the Reliability Models in the analysis significantly reduced the rate of wrong assignments. The results of the Reliability models for all taxonomic levels are summarised in Table 1 and Supplementary Table S5.

### Forestax tool evaluation on shotgun metagenomics data

Once we had validated the RecA protein sequence as a marker for bacterial taxonomy, we developed a tool called Forestax, which takes as inputs either paired-end raw reads, (meta)genomic assemblies or protein sequences, extract putative RecA proteins and provide a taxonomic assignment using the Taxa Assignment Models and Reliability Models. If raw reads are available, the tool can also infer relative abundances using the mean read coverage. Each step performed by the tool is described more thoroughly in the Methods (section “Forestax overview and evaluation on mock community samples”). To assess whether our approach could be used on shotgun metagenomics data, we used the tool to define bacterial relative abundances for 22 mock community samples. These samples were included in a recent paper evaluating the performance of taxonomy classification pipelines^30^, thus we compared our results to a set of commonly used pipelines using the same metrics (Sensitivity, False Positive Relative Abundance and Aitchison Distance).

Aitchison distances (AD) were used to assess the tool’s ability to predict relative abundances. At the genus level, AD values ranged from 8.26 (on the Bmock12 sample) to 21.72 (on the NIST-EG sample), while at the species level, values ranged from 10.24 (NIST-MIX-A) to 24.34 (Tourlosse).

False Positive Relative Abundances (FPRA) were used to quantify the wrong taxonomic assignments. At the genus level, FPRA values ranged from 0% (Bmok12, NIST-MIX-A, NIST-MIX-D, Tourlosse) to 28.33% (NIST-MIX-C); at the species level, values ranged from 0% (NIST-MIX-A, NIST-MIX-B, NIST-MIX-D, Amis-HiLo) to 42.84% (Bmock12).

Sensitivity values were used to quantify the right taxonomic assignments; at the genus level values ranged from 89.47% (Tourlousse) to 27.27% (NIST-MIX-A,NIST-MIX-B); at the species level, values ranged from 57.89% (Tourlousse) to 18.18% (NIST-MIX-C).

As shown in Figure 3, AD, FPRA and sensitivity values are coherent with other pipelines, especially when low-frequency taxa (with relative abundance < 2%) were not taken into account. All metrics are available in Supplementary Table S6.

**Figure 3.**
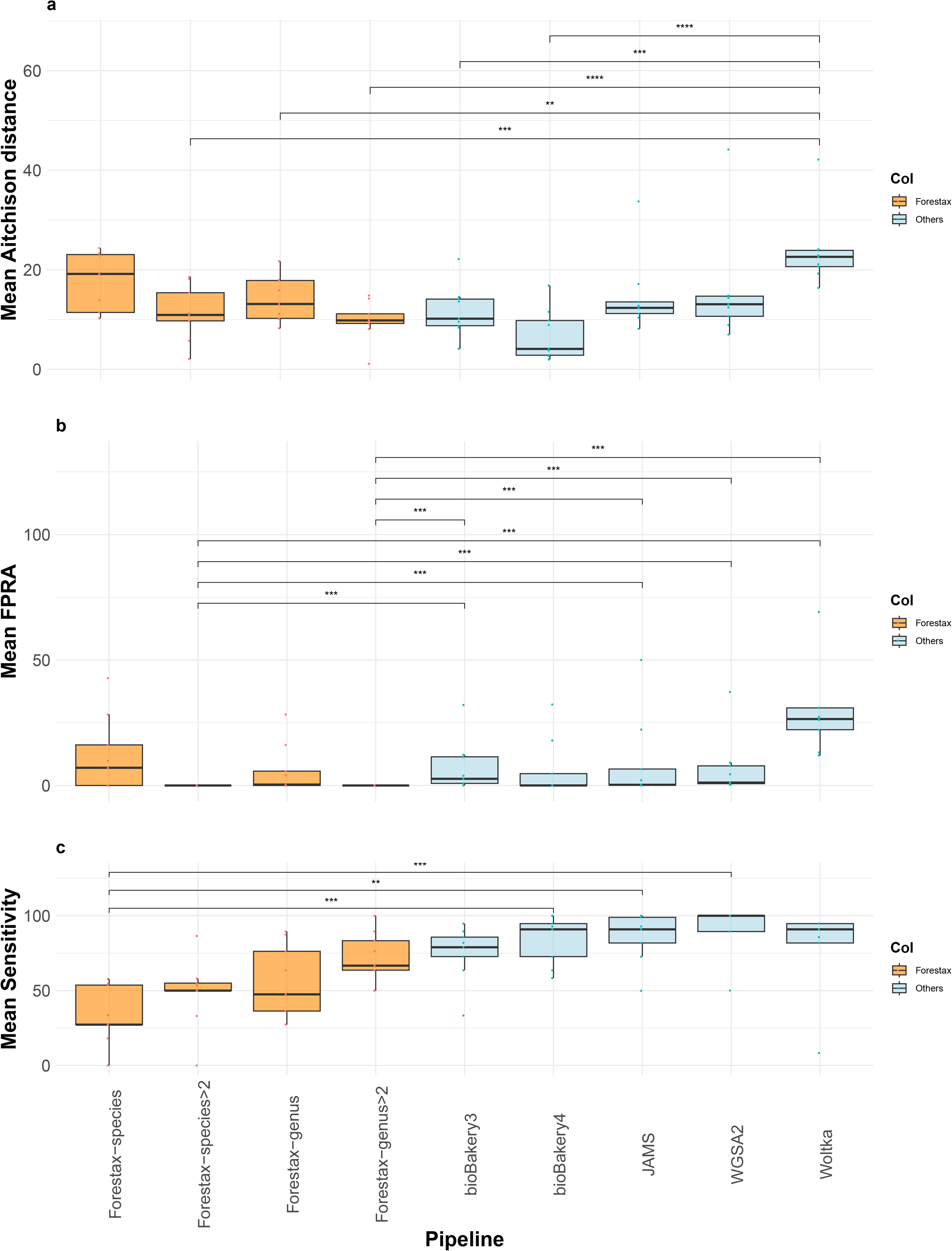
a) Boxplots showing the mean Aitchison distances between the true relative abundances in simulated microbial communities and the relative abundances estimated by different pipelines. The metrics for Forestax were computed at genus and species level, also removing low-abundance taxa (<2%) from the analysis. Statistically significant differences (Mann-Whitney U test, p < 0.05) are indicated by an asterisk (*). b) Boxplots showing the mean False Positive Relative Abundance (sum of all false positive relative abundances) estimated for simulated microbial communities by different pipelines. The metrics for Forestax were computed at genus and species level, also removing low-abundance taxa (<2%) from the analysis. Statistically significant differences (Mann-Whitney U test, p < 0.05) are indicated by an asterisk (*). c) Boxplots showing the mean Sensitivity (percentage of true positive species) for simulated microbial communities by different pipelines. The metrics for Forestax were computed at genus and species level, also removing low-abundance taxa (<2%) from the analysis. Statistically significant differences (Mann-Whitney U test, p < 0.05) are indicated by an asterisk (*).

## Discussion

The 16S rRNA gene (16S) is the most commonly used marker to define bacterial taxonomy. Despite being an effective and widely used marker, recent studies based on shotgun metagenomic sequencing have observed that several microbial groups evade 16S surveys^8,9^. These groups could constitute up to 25% of microbial diversity ^11^, with a main contributor being the Candidate Phyla Radiation (CPR)^12^. Moreover, the 16S is often found in multiple copies in the bacterial genome^31–33^ and can often be misassembled in whole genome sequencing (WGS) data.

For this reason, the first aim of this study was to identify a valid alternative genetic marker to identify and quantify the main groups of the Candidate Phyla Radiation. Performing orthology and phylogenetic analyses on a genomic dataset of CPR genomes, we identified the RecA protein as a valid target. RecA is present in almost all CPR strains, mostly found in single copy and phylogenetically coherent with the CPR evolutionary history: these features suggest that the protein is conserved within the CPR and the gene is not affected by horizontal gene transfer. These features are coherent with the fact that RecA is an essential enzyme involved in core cellular functions such as DNA repair and homologous recombination^34^.

Starting from this result and following up on other studies proposing RecA/*rec*A as a suitable taxonomy marker for specific bacterial groups ^35–39^, we expanded our analysis to all bacterial diversity. We created a pan-bacterial database of ∼300,000 RecA protein sequences and trained machine-learning models to predict the bacterial taxon on the basis of the sequence amino acid composition (“Taxa Assignment Models”). The models produced accurate predictions, indicating that RecA is indeed a valid taxonomic marker across the bacterial tree of life, able to distinguish bacterial genera and species.

There are two major drawbacks in using a single target and machine-learning for taxonomic assignment. Indeed, assignments could be compromised if the target sequence is incomplete/misassembled or if it belongs to a category (taxon) that is not known by the classifier. To deal with these issues, we trained additional models to assign a reliability score for each assignment (“Reliability Models”). Combining Taxa Assignment Models and Reliability Models allowed us to tackle these limits by placing sequences with low assignment reliability in the “Unclassified” category.

We implemented these models in a tool called Forestax, which, starting from reads, assemblies, overall protein content or already selected RecA protein sequences, automatically detects and/or estimates relative abundances of bacterial taxa.

Finally, the analysis of 22 mock communities with Forestax tool produced very few wrong assignments (i.e. low False Positive Relative Abundances) and estimated the bacterial relative abundances with metrics comparable to other commonly used pipelines based on whole genomic content or several target genes. The results were particularly robust for taxa with frequencies > 2% in the sample.

The 16S rRNA has been proposed as a marker gene before the genomic era. Despite the incredible success of its application, some limits have emerged with time. In this study we highlight that RecA is a powerful taxonomic marker in bacteria and we produced a fast tool suitable for screenings and large studies. Here, we show that the RecA protein sequence is a reliable marker for characterising bacteria up to the genus level, but uncertainties at the species level could be resolved using the nucleotide sequences. Moreover, being a single-copy conserved marker, the use of RecA could be suitable both for large shotgun metagenomics studies and for amplicon-based metagenomics. In the future, RecA could be a valid alternative marker to the 16S, combining a high sensitivity with low laboratory/informatic difficulties.

## References

1. Pérez-Cobas, A. E., Gomez-Valero, L. & Buchrieser, C. Metagenomic approaches in microbial ecology: an update on whole-genome and marker gene sequencing analyses. Microbial Genomics 6, (2020).

2. Laudadio, I. et al. Quantitative Assessment of Shotgun Metagenomics and 16S rDNA Amplicon Sequencing in the Study of Human Gut Microbiome. OMICS 22, (2018).

3. Tessler, M. et al. Large-scale differences in microbial biodiversity discovery between 16S amplicon and shotgun sequencing. Sci. Rep. 7, 1–14 (2017).

4. Rausch, P. et al. Comparative analysis of amplicon and metagenomic sequencing methods reveals key features in the evolution of animal metaorganisms. Microbiome 7, 1–19 (2019).

5. Liu, Y.-X. et al. A practical guide to amplicon and metagenomic analysis of microbiome data. Protein Cell 12, 315–330 (2020).

6. Woese, C. R., Kandler, O. & Wheelis, M. L. Towards a natural system of organisms: proposal for the domains Archaea, Bacteria, and Eucarya. Proc. Natl. Acad. Sci. U. S. A. 87, 4576 (1990).

7. Hugerth, L. W. & Andersson, A. F. Analysing Microbial Community Composition through Amplicon Sequencing: From Sampling to Hypothesis Testing. Front. Microbiol. 8, 274218 (2017).

8. Eloe-Fadrosh, E. A., Ivanova, N. N., Woyke, T. & Kyrpides, N. C. Metagenomics uncovers gaps in amplicon-based detection of microbial diversity. Nature Microbiology 1, 1–4 (2016).

9. Brown, C. T. et al. Unusual biology across a group comprising more than 15% of domain Bacteria. Nature 523, 208–211 (2015).

10. Maatouk, M., Rolain, J. M. & Bittar, F. Using Genomics to Decipher the Enigmatic Properties and Survival Adaptation of Candidate Phyla Radiation. Microorganisms 11, (2023).

11. Naud, S. et al. Candidate Phyla Radiation, an Underappreciated Division of the Human Microbiome, and Its Impact on Health and Disease. Clin. Microbiol. Rev. 35, (2022).

12. Castelle, C. J. et al. Biosynthetic capacity, metabolic variety and unusual biology in the CPR and DPANN radiations. Nat. Rev. Microbiol. 16, 629–645 (2018).

13. Starke, R., Pylro, V. S. & Morais, D. K. 16S rRNA Gene Copy Number Normalization Does Not Provide More Reliable Conclusions in Metataxonomic Surveys. Microb. Ecol. 81, 535–539 (2020).

14. Quince, C., Walker, A. W., Simpson, J. T., Loman, N. J. & Segata, N. Shotgun metagenomics, from sampling to analysis. Nat. Biotechnol. 35, 833–844 (2017).

15. Haft, D. H. & Tovchigrechko, A. High-speed microbial community profiling. Nat. Methods 9, 793–794 (2012).

16. Segata, N. et al. Metagenomic microbial community profiling using unique clade-specific marker genes. Nat. Methods 9, (2012).

17. Wood, D. E. & Salzberg, S. L. Kraken: ultrafast metagenomic sequence classification using exact alignments. Genome Biol. 15, (2014).

18. Koslicki, D. et al. ARK: Aggregation of Reads by K-Means for Estimation of Bacterial Community Composition. PLoS One 10, (2015).

19. Sunagawa, S. et al. Metagenomic species profiling using universal phylogenetic marker genes. Nat. Methods 10, (2013).

20. Parks, D. H. et al. GTDB: an ongoing census of bacterial and archaeal diversity through a phylogenetically consistent, rank normalized and complete genome-based taxonomy. Nucleic Acids Res. 50, D785–D794 (2021).

21. Olson, R. D. et al. Introducing the Bacterial and Viral Bioinformatics Resource Center (BV-BRC): a resource combining PATRIC, IRD and ViPR. Nucleic Acids Res. 51, (2023).

22. Hyatt, D. et al. Prodigal: prokaryotic gene recognition and translation initiation site identification. BMC Bioinformatics 11, 1–11 (2010).

23. Buchfink, B., Reuter, K. & Drost, H.-G. Sensitive protein alignments at tree-of-life scale using DIAMOND. Nat. Methods 18, 366–368 (2021).

24. Galperin, M. Y. et al. COG database update: focus on microbial diversity, model organisms, and widespread pathogens. Nucleic Acids Res. 49, D274–D281 (2020).

25. Edgar, R. C. MUSCLE: a multiple sequence alignment method with reduced time and space complexity. BMC Bioinformatics 5, 1–19 (2004).

26. Price, M. N., Dehal, P. S. & Arkin, A. P. FastTree 2 – Approximately Maximum-Likelihood Trees for Large Alignments. PLoS One 5, e9490 (2010).

27. Prjibelski, A., Antipov, D., Meleshko, D., Lapidus, A. & Korobeynikov, A. Using SPAdes De Novo Assembler. Curr. Protoc. Bioinformatics 70, (2020).

28. Langmead, B. & Salzberg, S. L. Fast gapped-read alignment with Bowtie 2. Nat. Methods 9, 357.

29. Danecek, P. et al. Twelve years of SAMtools and BCFtools. Gigascience 10, giab008 (2021).

30. Valencia, E. M., Maki, K. A., Dootz, J. N. & Barb, J. J. Mock community taxonomic classification performance of publicly available shotgun metagenomics pipelines. Scientific Data 11, 1–24 (2024).

31. Sun, D. L., Jiang, X., Wu, Q. L. & Zhou, N. Y. Intragenomic heterogeneity of 16S rRNA genes causes overestimation of prokaryotic diversity. Appl. Environ. Microbiol. 79, (2013).

32. Pei, A. Y. et al. Diversity of 16S rRNA genes within individual prokaryotic genomes. Appl. Environ. Microbiol. 76, (2010).

33. Ibal, J. C., Pham, H. Q., Park, C. E. & Shin, J.-H. Information about variations in multiple copies of bacterial 16S rRNA genes may aid in species identification. PLoS One 14, (2019).

34. Karlin, S. & Brocchieri, L. Evolutionary conservation of RecA genes in relation to protein structure and function. J. Bacteriol. 178, 1881 (1996).

35. Eisen, J. A. The RecA Protein as a Model Molecule for Molecular Systematic Studies of Bacteria: Comparison of Trees of RecAs and 16S rRNAs from the Same Species. J. Mol. Evol. 41, 1105 (1995).

36. Thompson, C. C. et al. Use of recA as an alternative phylogenetic marker in the family Vibrionaceae. Int. J. Syst. Evol. Microbiol. 54, 919–924 (2004).

37. Liu, W., Li, L., Khan, M. A. & Zhu, F. Popular molecular markers in bacteria. Mol. Gen. Microbiol. Virol. 27, 103–107 (2012).

38. Rossi, F., Dellaglio, F. & Torriani, S. Evaluation of recA gene as a phylogenetic marker in the classification of dairy propionibacteria. Syst. Appl. Microbiol. 29, (2006).

39. Meyers, P. R. Analysis of recombinase A (recA/RecA) in the actinobacterial family Streptosporangiaceae and identification of molecular signatures. Syst. Appl. Microbiol. 38, (2015).

