## Supplementary figures for "RecA is a reliable marker for bacterial taxonomy, even in the Candidate Phyla Radiation"

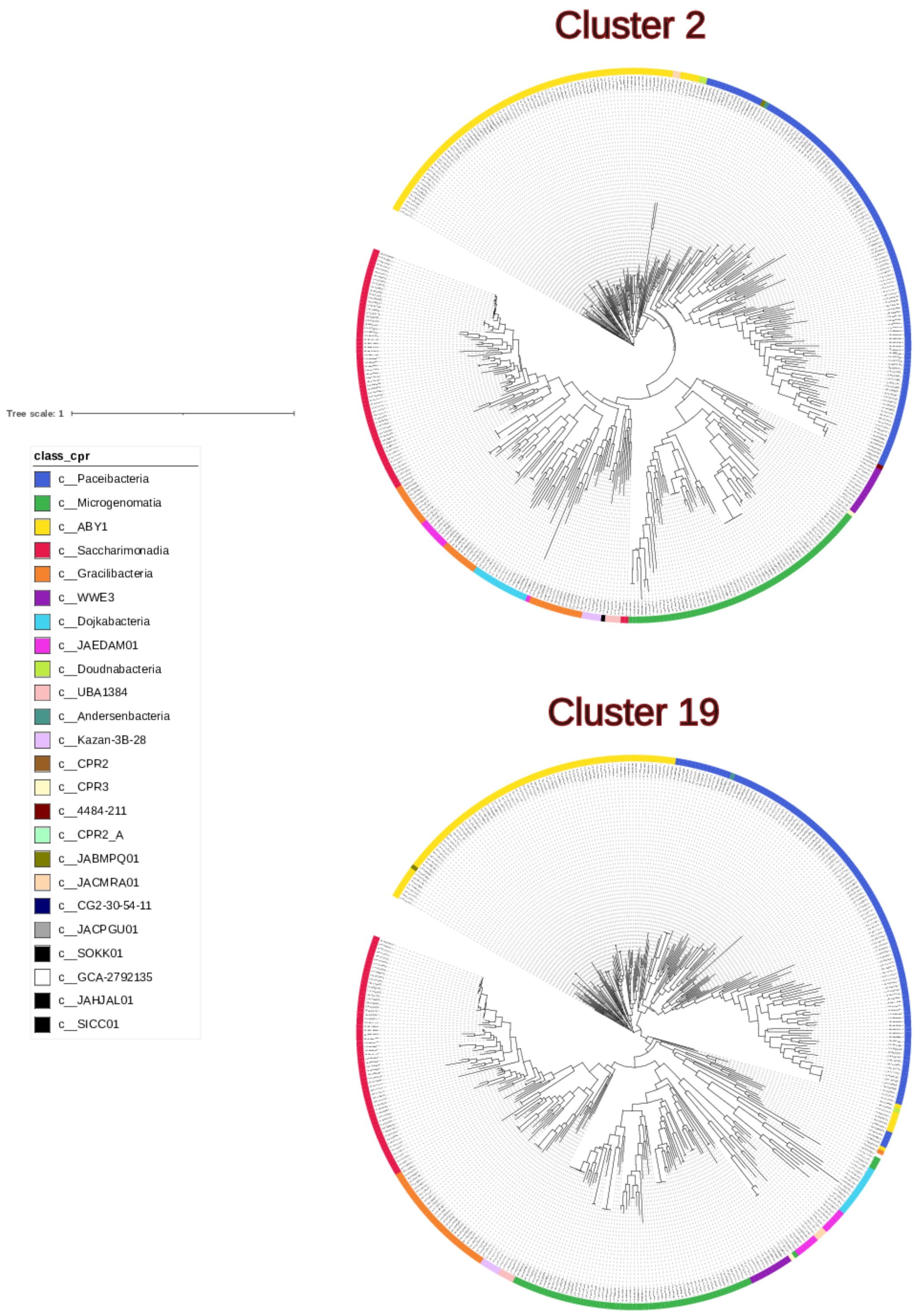


**Figure S1.** ML Phylogenetic trees obtained from two candidate protein markers within a refined dataset of Candidate Phyla Radiation (CPR) genomes

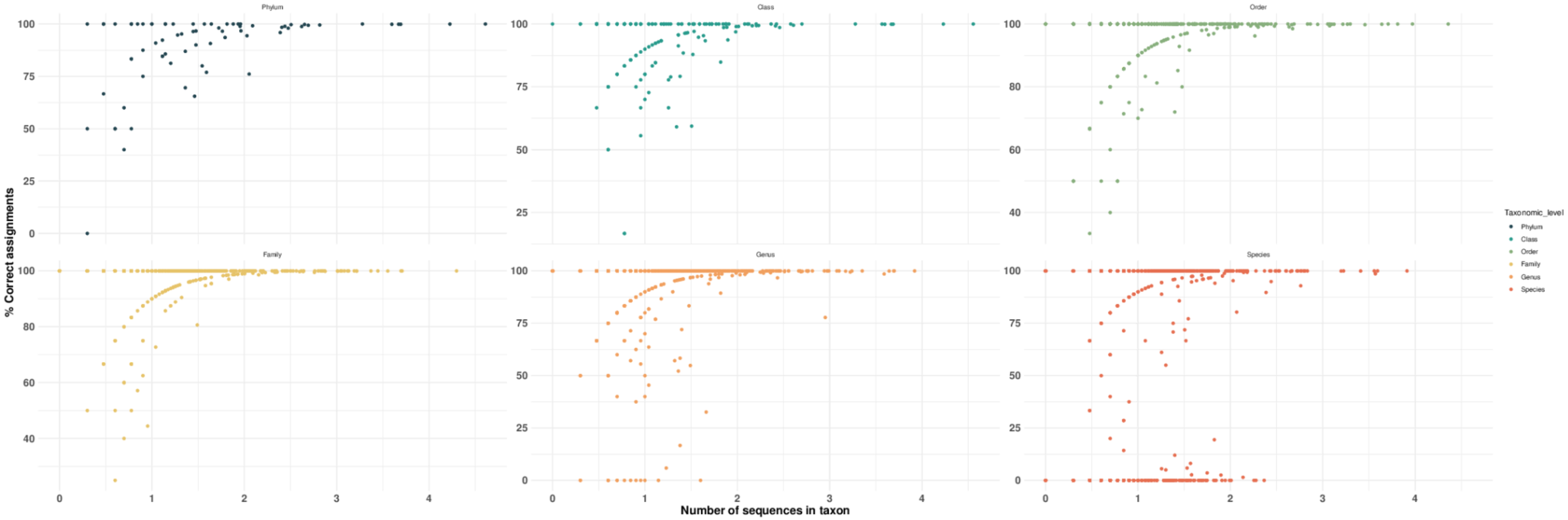


**Figure S2.** Plot showing the relationship between the logarithm of the number of RecA sequences available from each bacterial taxon and the percentage of sequences correctly assigned to the taxon by the Taxa Assignment models.

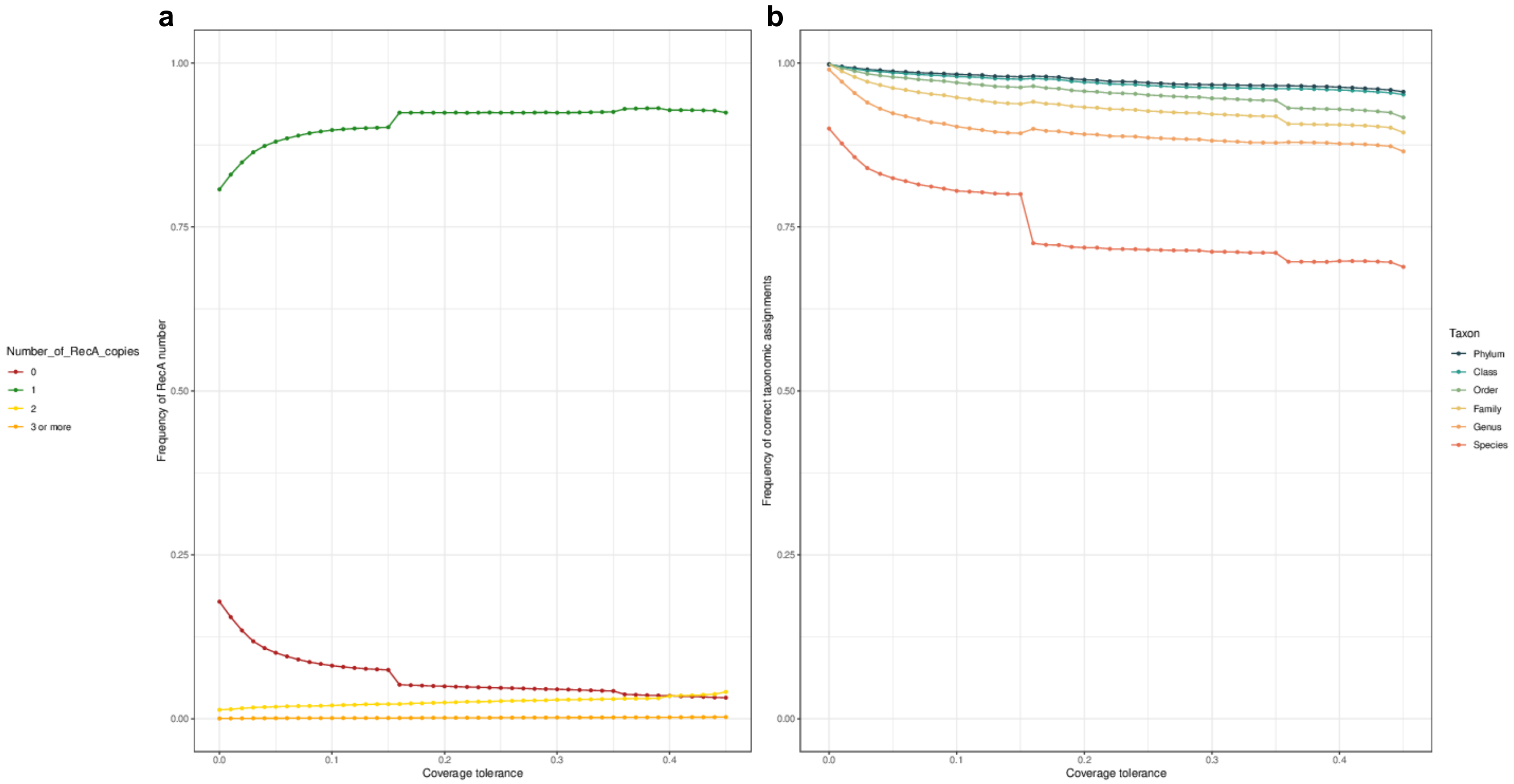


**Figure S3.** a) Relationship between the coverage tolerance (i.e. the inclusion of sequences annotated against the RecA sequences in the COG database with a coverage progressively above or below 1, see Methods) and the frequency of different copy numbers of RecA obtained from the genomes. b) Relationship between the coverage tolerance and the frequency of correct taxonomic assignments for each taxonomic level

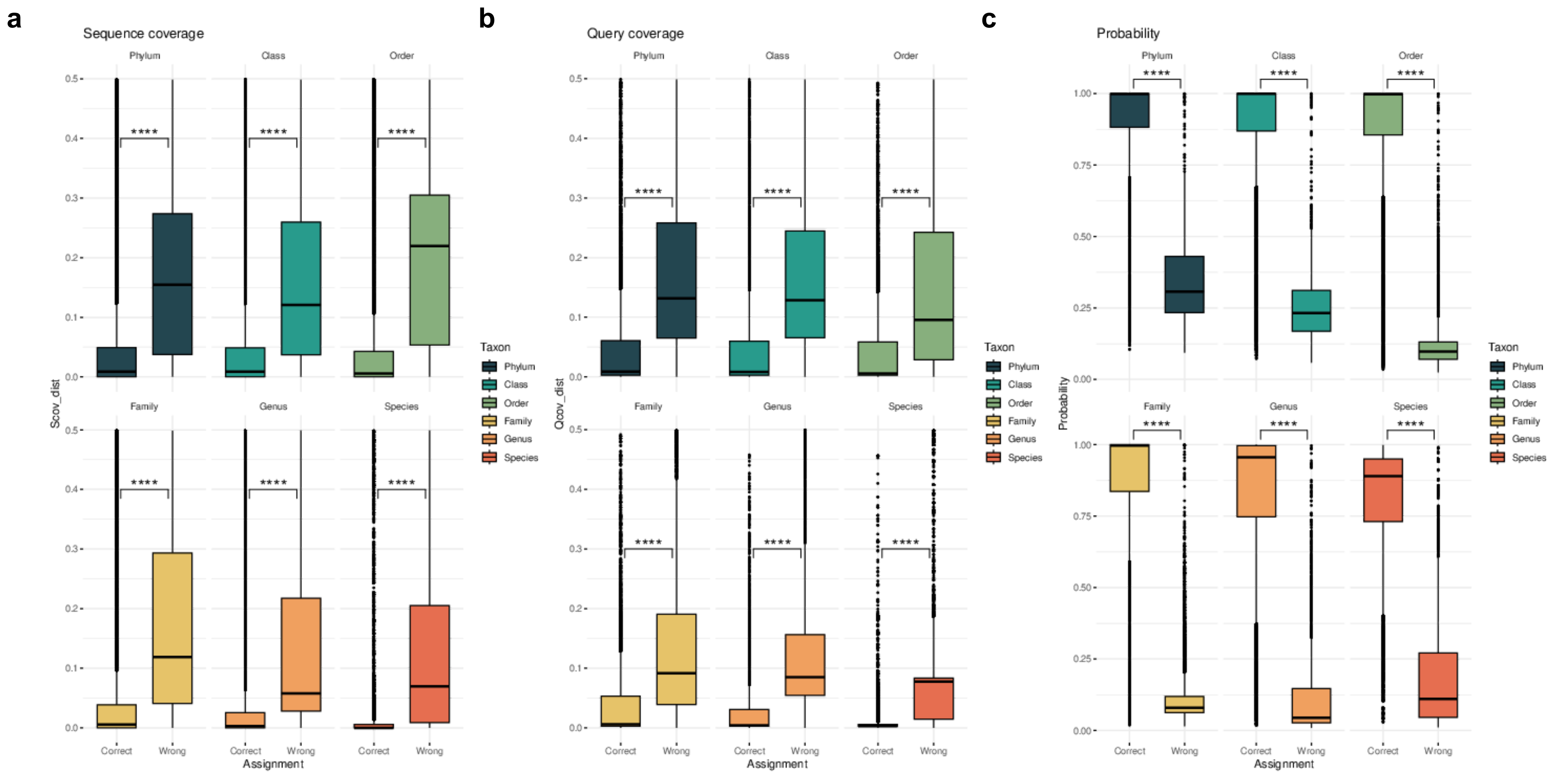


**Figure S4.** a) Boxplots showing the sequence coverage distance (Scov_dist, see Methods) for RecA sequences assigned to the correct or wrong at each taxonomic level. b) Boxplots showing the query coverage distance (Scov_dist, see Methods) for RecA sequences assigned to the correct or wrong at each taxonomic level. c) Boxplots showing the assignment probability for RecA sequences assigned to the correct or wrong at each taxonomic level. Statistically significant differences (Mann-Whitney U test) are indicated by the asterisks (*: p <0.05; ** **:** p < 0.01; *** : p < 0.001; **** : p < 0.0001).
